# SNRNP70 interacts with TDP-43 to promote RNP granule localisation and regulate motor neuron development

**DOI:** 10.64898/2026.07.17.738863

**Authors:** Tilly Baldacchino, Joshua Lloyd-Jones, Chloe M. Edwards, Stephanie M.E. Jones, James Ellams, Johanna Ganssauge, Corin Liddle, Akshay Bhinge, Nikolas Nikolaou

## Abstract

SNRNP70 is a core spliceosomal protein that localises to both the nucleus and cytoplasm. Previous studies have implicated SNRNP70 in regulating axonal stability and the transport of specific mRNAs during motor neuron development in zebrafish. Although the molecular functions and protein interactions of SNRNP70 in pre-mRNA splicing are well established, the mechanisms underlying its cytoplasmic functions remain poorly understood. Here, we show that SNRNP70 and TDP-43 exhibit similar localisation patterns in developing and mature neurons and co-associate in both nuclear and non-nuclear compartments, including axonal projections. We identify a functional interaction between SNRNP70 and TDP-43 that is essential for motor neuron development and demonstrate that the recruitment of SNRNP70 to cytoplasmic ribonucleoprotein (RNP) granules depends on TDP-43. These findings identify a previously unrecognised cytoplasmic function of TDP-43 in directing SNRNP70-containing RNP granule assembly, thereby linking TDP-43 to the splicing-independent functions of SNRNP70 during motor neuron development.

## Introduction

While most mRNA processing (e.g., 5’ capping, splicing, and 3’ polyadenylation) occurs in the nucleus, extra-nuclear post-transcriptional events are also critical for controlling mRNA fate and gene expression. Events such as mRNA transport, localisation, local translation and stability/degradation play a critical role in regulating how much, when, and where proteins are synthesised. This is especially evident in neurons, where the soma harbouring the mRNA-producing nucleus is separated from the final destination and translation point of many mRNA transcripts by millimetres or even centimetres. This underscores the importance of mRNA processing at synapses, growth cones and dendritic and axonal branch points for normal neuronal development and function (Taylor & Nikolaou, 2024). The extra-nuclear processing, transport, translation and eventual degradation of mRNAs is regulated by several RNA-binding proteins (RBPs). These proteins play crucial roles in controlling the fate of the mRNA and how it is utilised within the cell (Darnell & Richter, 2012; Street et al., 2024).

SNRNP70 (U1-70K) plays a key role in mRNA splicing in the nucleus. It is a component of the U1 small nuclear ribonucleoprotein (snRNP) complex, which forms part of the spliceosome, the molecular machine responsible for removing introns from precursor mRNA (pre-mRNA) during the process of splicing (Will & Lührmann, 2011). SNRNP70 interacts with the other two U1-specific proteins (U1-A and U1-C), the core components of the spliceosome as well as with U1 snRNA, the RNA component of the U1 snRNP complex(Kondo et al., 2015). The U1 snRNA is essential for recognizing the 5’ splice site of pre-mRNA during the splicing process. SNRNP70 helps stabilise the interaction between U1 snRNA and the pre-mRNA, which is critical for initiating the splicing reaction (Kondo et al., 2015; Pomeranz Krummel et al., 2009).

Besides its well-characterised role in mRNA splicing, SNRNP70 colocalises with its mRNA targets in cytoplasmic regions (Lee et al., 2025; Nikolaou et al., 2022). We have recently demonstrated that the cytoplasmic pool of SNRNP70 is essential for motor neuronal connectivity in zebrafish (Nikolaou et al., 2022). At the transcriptome level, SNRNP70 was found to regulate the stability and transport of cytoplasmic mRNAs in axons (Nikolaou et al., 2022). However, the molecular mechanisms by which extra-nuclear SNRNP70 regulates neuronal connectivity are still unclear.

One possible scenario is that SNRNP70 directly interacts with mRNA molecules to regulate their cytoplasmic processing. Although SNRNP70 has its own RNA recognition motif (RRM) that enables interactions with the U1 snRNA (Rossi et al., 1996) and the potential nuclear enrichment of transcripts (Lubelsky et al., 2021), there is little evidence currently to suggest that SNRNP70 can directly bind to mRNAs outside the nucleus. An alternative hypothesis is that SNRNP70 regulates extra-nuclear mRNA processing indirectly through associations with other RBPs. Indeed, evidence suggests that besides the U1 snRNP complex, SNRNP70 can associate with several RBPs, including Survival Motor Neuron (SMN) (Stejskalová & Staněk, 2014) and Gemin5 (So et al., 2016) as well as mRNA processing factors and ribosomal subunits (Bishof et al., 2018).

A well-known mRNA processing regulator is TAR DNA-binding protein 43 (TDP-43). TDP-43 is a ubiquitously expressed RBP that predominantly localises to the nucleus, where it has fundamental RNA-processing activities including RNA transcription (Morera et al., 2019) and alternative splicing (Ma et al., 2022). It comprises of an N-terminal domain (NTD), two tandem RNA recognition motifs (RRMs; RRM1 and RRM2), and an intrinsically disordered C-terminal domain (CTD). Previous studies have shown that the RRMs of TDP-43 mediate its binding to GU-rich repeat regions within transcripts (Bhardwaj et al., 2013; Kuo et al., 2009, Kuo et al., 2014), which are often localised in the introns and 3’ UTRs of its targets. RNA binding facilitates the high-affinity multimerisation of TDP-43 (Carlos Rengifo-Gonzalez et al., 2021), which is thought to enable the appropriate splicing of its target introns.

Interestingly, TDP-43 shuttles between the nucleus and the cytoplasm and participates in various cytoplasmic mRNA processing events including the control of mRNA stability and abundance, ribonucleoprotein (RNP) granule formation and transport, local mRNA translation, and stress granule dynamics (reviewed by García Morato et al., 2023). It is thought that TDP-43 acts as an RNP granule protein within actively transported RNP granules. Research by Nagano and colleagues (2020) demonstrated a role for TDP-43 in neuronal granule transport and in the regulation of protein synthesis in axons. By depleting TDP-43 within neurons, they observed a decrease in expression of ribosomal protein mRNAs, along with an overall decrease in protein synthesis (Nagano et al., 2020). Additional research by Briese and colleagues (2020) also demonstrated that depletion of TDP-43 results in deficits in axonal outgrowth, concomitant with alterations in the axonal and somatodendritic transcriptomes (Briese et al., 2020). Overall, these recent studies suggest that TDP-43 plays a significant role in regulating mRNA transport and facilitating local protein synthesis within neuronal projections and terminals.

In this study, we examined whether the functions of SNRNP70 in mRNA processing outside the nucleus and in motor connectivity rely on its association with Tardbp, which is the native TDP-43 protein in zebrafish. Our results indicate that SNRNP70 and Tardbp interact within cells, with many of these co-associations mapping to both cytoplasmic and axonal regions of neurons. Moreover, we show that these two proteins function synergistically during motor neuron development. Finally, we provide evidence that Tardbp is essential for the recruitment and localisation of SNRNP70 to RNP granules, thereby enabling SNRNP70-mediated mRNA processing. Our findings provide new insights into the molecular mechanisms by which SNRNP70 regulates the transcriptome in neuronal projections.

## Materials and methods

### Animals

Zebrafish were reared at 28.5°C on a 14-hours light/10-hours dark cycle. Embryos produced by natural crosses were raised in E3 media (5 mM NaCl, 0.17 mM KCl, 0.33 mM CaCl_2_, 0.33 mM MgSO_4_). We used the following existing transgenic and mutant lines: *snrnp70^kg163^ (Nikolaou et al., 2022)*, *TgKI(snrnp70-eGFP)^ba9^* (Lloyd-Jones et al., in preparation); *Tg(ubi:ERT2-Gal4)^nim10Tg^ (Gerety et al., 2013)*, *Tg(UAS:hSNRNP70-eGFP)^kg323Tg^* (Nikolaou et al., 2022), *Tg(huC:GFP)* (Park et al., 2000). This work was approved by the local Animal Welfare and Ethical Review Bodies (University of Bath and University of Exeter) and was carried out in accordance with the Animals (Scientific Procedures) Act, 1986, under project license (PP8698401) from the United Kingdom Home Office.

### Generation of F0 *tardbp* crispant model

For the generation of F0 *tardbp* crispants, a CRISPR/Cas9 strategy was used with three guides targeting the *tardbp* locus. The three CRISPR RNAs (crRNAs) were designed using the CRISPR-Cas9 guide RNA (gRNA) design checker (Integrated Design Technologies) and CHOPCHOP algorithm (Labun et al., 2019). As previously described (Kroll et al., 2021), equimolar concentration of individual crRNAs and tracrRNA were combined and diluted to 61 μM using Nuclease Duplex Buffer (Integrated DNA Technologies). Samples were then heated to 95°C for 5 minutes and then cooled on ice for 2 minutes. Mock injection mixtures lacked the *tardbp* targeting crRNAs. Individual gRNAs and Alt-R S.p. Cas9 Nuclease V3 (Integrated DNA Technologies) were combined to a 1:1 ratio and heated to 37°C for 5 minutes to form RNP complexes. The three RNPs were pooled together in equal volumes to a final concentration of 30.5 µM. 1nl was injected into the yolk of embryos at one-cell stage.

### Validation of F0 *tardbp* crispant

To confirm the partial deletion of the *tardbp* locus was successful, DNA from individual embryos was extracted from uninjected and *tardbp* crispant embryos by alkali lysis and analysed by PCR. PCR reactions were assembled using DreamTaq Green Master Mix (Thermofisher Scientific-K1081) with the addition of 1M betaine (Merck- B0300) and run on an Eppendorf MasterCycler using the following thermocycling conditions: initial denaturation at 95°C for 3 minutes, 35 cycles of [95°C for 30 seconds, 63°C for 30 seconds, 72°C for 1 minute per kb] then a final extension of 72°C for 10 minutes. The following primers were used for PCR reactions: *tardbp* forward 5’- CGGATCTCACGACCAATCTAAG -3’ *tardbp* reverse *5’- TGACTTCCCCAAATGTACCGA -*3’. Western blotting as described below was performed to confirm reduction of the protein using the TDP-43 antibody (Proteintech-12892-1-AP; 1:500 dilution).

### 4-OHT treatment for Gal4 induction

4-hydroxytamoxifen (4-OHT) was dissolved in ethanol at a final stock concentration of 50 mM and stored in the dark at –20°C. To induce Gal4 activity in *Tg(ubi:ERT2-Gal4);Tg(UAS:hSNRNP70-eGFP)* offspring, embryos at sphere stage (4 hpf) were incubated with E3 media containing 5 μM of 4-OHT in the dark at 28.5°C. They remained in 4-OHT-containing E3 media until required for subcellular fractionation (96 hpf), immunohistochemistry (96 hpf) or primary neuron culture (24 hpf).

### Primary neuron culture

Embryos from *Tg(ubi:ERT2-Gal4);Tg(UAS:hSNRNP70-eGFP)* or *TgKI(snrnp70-eGFP)* lines were collected and incubated at 28.5°C overnight in E3 medium containing 1X Penicillin-Streptomycin and 50 μg/mL gentamicin. Additionally, E3 medium contained 5 μM 4-OHT to induce Gal4 activity where relevant. At 24 hpf, embryos expressing eGFP were selected and used for culture based upon methods published by (Taylor and Houart, 2024). The following modifications were made: dissociation of cells from zebrafish tissue was undertaken at room temperature with agitation via pipetting with a P1000 for a maximum of 25 minutes; coverslips decontaminated with 70% EtOH were incubated in a 24 well plate with 0.1 mg/mL poly-D-lysine at 4°C overnight, followed by 2 hours incubation with 0.5 μg/mL laminin in HBSS; cells were seeded at a density of 300-400 embryos per mL by pipetting 50 μL cell suspension onto coated coverslips; 1 mL of zebrafish neural media was added after 90 minutes of incubation, and a further 1mL was added at 6 days *in vitro* (div). For inducing Gal4 activity in *Tg(ubi:ERT2-Gal4);Tg(UAS:hSNRNP70-eGFP)*-derived primary neurons, 0.5 μM of 4-OHT was included to the media. In mock-treated cultures, the equivalent amount of carrier (i.e., ethanol) was used instead. Cultures were kept at room temperature in the dark. Cells were fixed at 12-div with 4% PFA for 10 minutes at room temperature followed by 3 washes with PBS and incubation with 100% methanol for 10 minutes at room temperature. Coverslips were washed with PBS and stored at 4°C to be used for immunocytochemistry or proximity ligation assay.

### Human iPSC-derived motor neurons

Motor neurons were generated from TDP-43-GFP knock-in iPSCs as described previously (Ganssauge et al., 2025).

### Immunohistochemistry (IHC)

Whole-mount immunohistochemistry was performed as previously described (Nikolaou et al., 2022). Briefly, 4% paraformaldehyde-fixed embryos were washed with PBS, permeabilised with 0.25% Trypsin in PBS for 10 minutes and blocked with 10% goat serum/PBS at room temperature (RT) for 1 hour. Embryos were incubated with mouse anti-SV2 (DSHB- AB2315387; 1:100 dilution) in 10% goat serum/PBST (PBS + 1%Triton X-100) at 4°C overnight, washed with PBST and incubated at 4°C overnight with Alexa Fluor 488 secondary antibody diluted in 10% goat serum/PBST at 1:500. Samples were mounted on glass slides and imaged using Leica TCS SP8 confocal microscope equipped with spectral detectors and a 40x/1.3 NA oil objective. Excitation was 488 nm. Images were captured at 0.28 x 0.28 μm resolution (1024 x 1024 pixels) and 1 A.U. Cryosections were rehydrated in PBS for 10 minutes and then blocked in for 1 hour before incubating with chick anti-GFP (Fisher Scientific-PA19533; 1:500 dilution), rabbit anti-TDP-43 (Proteintech-12892-1-AP; 1:400 dilution) and mouse anti-acetylated α-Tubulin (Sigma-T7451; 1:500 dilution) at 4°C overnight. Sections were then washed and incubated with Alexa Fluor secondary antibodies (goat anti-Chick-488, goat anti-rabbit-568, goat anti-mouse-633) diluted in 1% goat serum in PBST at 1:1,000 for 2 hours at room temperature. Sections were incubated with 1.25 μg/mL DAPI for 1 minute, washed in PBS and covered with glass slides using FluorSave Reagent (Merck- 345789). Samples were imaged using Olympus BX63LF/FV3000 confocal microscope equipped with spectral detectors and a 100x/1.25 NA oil objective. Excitation was 405 nm (for DAPI), 488 nm (for GFP), 568 nm (for Tardbp) and 633 nm (for acetylated α-Tubulin). Images were captured at 0.2 x 0.2 μm resolution (1024 x 1024 pixels) and 1 A.U.

### Immunocytochemistry (ICC)

Fixed coverslips were blocked with 5% goat serum/PBST (PBS + 0.1% Triton X-100) for 30 minutes at room temperature. Following removal of blocking solution, coverslips were incubated with the following primary antibodies: chick anti-GFP (Fisher Scientific-PA19533; 1:500 dilution), mouse anti-acetylated α-Tubulin (Sigma-T7451; 1:500 dilution), rabbit anti-TDP-43 (Proteintech-12892-1-AP; 1:400 dilution) in 1% goat serum in PBST at 4°C overnight. Three washes in PBS were performed followed by incubation with Alexa Fluor secondary antibodies (goat anti-chick-488, goat anti-mouse-568, goat anti-rabbit-633) diluted in 1% goat serum in PBST at 1:1,000 for 1 hour at room temperature. After 3 washes in PBS, coverslips were incubated with 1.25 μg/mL DAPI for 1 minute before 2 further washes in PBS. Coverslips were mounted onto glass slides (cells facing down) with 10μl FluorSave Reagent (Merck- 345789). Samples were imaged using Olympus BX63LF/FV3000 confocal microscope equipped with spectral detectors and a 100x/1.25 NA oil objective. Excitation was 405 nm (for DAPI), 488 nm (for GFP), 568 nm (for acetylated α-Tubulin) and 633 nm (for Tardbp). Images were captured at 0.12 x 0.12 μm resolution (1024 x 1024 pixels) and 1 A.U.

### Proximity ligation assay (PLA)

Fixed coverslips were quenched with 50mM Na_4_Cl and then permeabilised with PBST (PBS + 0.2% Triton X-100) for 15 minutes each at room temperature. PLA was performed based upon the Duolink® PLA Fluorescence Protocol, using Duolink® In Situ PLA® Probe Anti-Rabbit PLUS (Merck-DUO92002), PLA® Probe Anti-Goat PLA MINUS (Merck-DUO92006), and Detection Reagents Orange (Merck-DUO92007). For zebrafish primary neurons, primary antibodies used were: rabbit anti-TDP-43 (Proteintech-12892-1-AP; 1:400 dilution) and goat anti-GFP (Abcam-ab6673; 1:400 dilution). For human iPSc-derived motor neurons, primary antibodies used were: rabbit anti-SNRNP70 (Sigma-AV40276; 1:200 dilution) and goat anti-GFP (Abcam-ab6673; 1:400 dilution). Following PLA, an ICC was completed as above using chick anti-GFP (Fisher Scientific-PA19533; 1:500 dilution), mouse anti-acetylated α-Tubulin (Sigma-T7451; 1:500 dilution) and mouse anti-Neurofilament (Merck-MAB1621; 1:500 dilution) primary antibodies, and Invitrogen Alexa Fluor goat anti-mouse 488, and donkey anti-chick 647 secondary antibodies. Samples were imaged using Leica TCS SP8 confocal microscope equipped with spectral detectors and a 100x/1.44 NA oil objective. Excitation was 405 nm (for DAPI), 488 nm (for acetylated α-Tubulin), 561 nm (for PLA) and 647 nm (for GFP). Images were captured at 0.05 x 0.05 μm resolution (1024 x 1024 pixels) and 1 A.U.

### PLA quantification

For analysing PLA, images were split by channel and z-stack before inputting into Cell Profiler (Stirling et al., 2021). The nucleus was defined first using the ‘IdentifyPrimaryObjects’ function and DAPI channel. Subsequently, the soma was defined using ‘IdentifySecondaryObjects’ function with the Ac α-Tub channel as ‘image’ and nucleus as ‘object’. To specify the soma cytoplasm, ‘IdentifyTertiaryObjects’ function was used by selecting soma as the larger ‘object’ and nucleus as the smaller ‘object’. For delineating axons, the ‘IdentifyPrimaryObjects’ function and the Ac α-Tub channel were first used, identifying Ac α-Tub as a primary ‘object’, before ‘IdentifyTertiaryObjects’ was used by selecting Ac α-Tub as the larger ‘object’ and soma as the smaller ‘object’. Once all cellular compartments were defined, the PLA channel was enhanced using the ‘EnhanceOrSupressFeatures’ function before using the ‘IdentifyPrimaryObjects’ function and the enhanced PLA image to specify PLA dots. Subsequently, PLA dots were counted in the following three compartments defined above: nucleus, soma cytoplasm, and axons. Finally, to calculate the surface area covered by each compartment, the ‘MeasureImageAreaOccupied’ was used to obtain a total pixel count before converting this into μm^2^. The sum of all PLA dots and surface area across all z-planes was obtained. The normalised number of PLA dots per μm^2^ was finally calculated for each compartment.

### Subcellular fractionation

Approximately 200 larvae at 4 dpf were anesthetised using MS222 and transferred into a Dounce tissue homogeniser with 1:5 weight/volume CLB buffer pH 8 (HEPES (10 mM), NaCl (10 mM), KH_2_PO4 (1.028 mM), NaHCO_3_ (5 mM), EDTA (5 mM), CaCl_2_ (1 mM) MgCl_2_ (1.05 mM) supplemented with protease, phosphatase and Granzyme B inhibitors) for homogenisation. Samples were deyolked by 10 strokes of a loose pestle A followed by a 5-minute centrifugation at 300xg. The pellet was resuspended in CLB and homogenised using 10 strokes with a tight pestle B. The osmotic balance was re-established with sucrose to a final concentration of 0.25 mM to form the total lysate. The cytoplasmic and nuclear compartments were separated via centrifugation at 6,300xg for 10 minutes and the supernatant was removed as the cytoplasmic compartment. The pellet was resuspended in 0,1% TSE buffer (Tris (1 mM), sucrose (30 mM), EDTA (0.1 mM), IGEPAL (0.01%) supplemented with protease, phosphatase, and Granzyme B inhibitors), centrifuged at 4,000xg for 5 minutes and the supernatant was discarded. This cycle was repeated 3-5 times until the supernatant ran clear. The nuclear pellet was finally resuspended in 1% TSE buffer, incubated on ice for 30 minutes, and sonicated for 15 seconds.

### Co-immunoprecipitation (Co-IP)

5 μg of rabbit anti-GFP (Abcam-ab6556) or rabbit IgG (Sigma-I5381) (control) antibodies were incubated with 50μL magnetic protein A dynabeads in 500 μl pre-binding buffer (NaCl (150 mM), Tris pH 8.0 (50 mM), IGEPAL (1%)) at 4°C for 4 hours. Equal amounts of total lysate were added following washing of the beads with IP wash buffer (NaCl (150 mM), Tris pH 8.0 (10 mM), IGEPAL (0.1%)) and incubated overnight at 4°C. The flowthrough was collected as the unbound fraction, beads were washed 3 times with IP wash buffer and then protein was eluted from the beads in a mixture containing 87.5 μL DTT and LDS at a 1:2.5 ratio at 70°C for 5 minutes. The protein was detected using western blotting as described below.

### Western blotting (WB)

Fractionated samples were prepared with 1X Bolt Sample Reducing agent (Thermo Fisher Scientific-B0009) and 1X Bolt LDS Sample Buffer (Thermo Fisher Scientific-B0007) and were denatured at 70 °C for 5 minutes. Western blotting was undertaken using 4-12% BisTris midi protein gels (Thermo Fisher Scientific-WG1401BOX) and transferred using the Invitrogen iBlot2 system to PVDF membranes (Thermo Fisher Scientific-IB24001). 5% milk in TBST (TBS + 0.1% Tween-20) was used for blocking and membranes were incubated with primary antibodies diluted in 1% milk in TBST overnight at 4°C. The following primary antibodies were used: goat anti-GFP (Abcam-ab6673; 1:1,000 dilution), rabbit anti-GFP (Abcam-ab6556; 1:500 dilution), mouse anti-GFP (Thermo Fisher Scientific-MA1-952; 1:1,000 dilution), rabbit anti-TDP-43 (Proteintech-12892-1-AP; 1:500 dilution), mouse anti-α-Tubulin (Cell Signaling Technology-3873; 1:5000 dilution), mouse anti-Lamin B1 (Proteintech-66095-1-Ig; 1:1,000 dilution) and mouse anti-Histone H3 (Cell Signaling Technology-96C10; 1:1,000 dilution). Following few quick washes for an hour, membranes were incubated with the following HRP-conjugated secondary antibodies, all diluted at 1:10,000 in 1% milk in PBST, at room temperature for 1 hour: goat anti-rabbit HRP (Abcam-ab205718), goat anti-mouse HRP (Abcam- ab205718), donkey anti-goat HRP (Abcam-ab6885). Following few quick washes for an hour, membranes were incubated with ECL substrate (Thermo Fisher Scientific-34580) and a Licor Odyssey Fc imager was used for imaging chemiluminescent signals.

### Analysis of motor neuron nerve length

Images were opened in Fiji and motor nerves in somites 9-12 were traced using the SNT (simple neurite tracer) plugin. Motor nerve length was measured from the spinal cord exit point to the most distal end of the nerve.

### Live imaging of RNP granules

*TgKI(snrnp70-eGFP)* fertilised eggs were injected with 100 μM Cy5-UTP solution alone or alongside gRNAs targeting *tardbp* gene (see above) at one-cell stage. Embryos at 52 hpf were mounted in 1% low melting point agarose (Thermo Fisher Scientific- 16520050). Imaging was performed using a Zeiss LSM 880 Fast Airyscan confocal microscope equipped with GaAsP spectral detectors and a 40x/1.2 N.A. water-immersion objective (Carl Zeiss). Excitation was provided using 488 nm (for GFP) and 633 nm (for Cy5) solid-state lasers. High-resolution images were captured at 2.6 Hz at a 0.04 x 0.04 μm resolution (488 x 488 pixels) before being processed for Airyscan imaging.

### RNP granule analysis

Raw image files were processed using the Airyscan processing workflow in ZEN prior to downstream analysis. Processed files were then imported into IMARIS v10.1.2 for image analysis. Background subtraction was applied consistently across all files before segmentation and quantification.

RNP granules were identified from the Cy5-UTP fluorescence signal. An initial segmentation threshold was optimised using representative images and then batch-applied consistently across comparable image files to minimise subjective adjustment between samples. Individual RNP granule regions of interest were then used to generate masks for downstream intensity and colocalisation measurements. This ensured that time-dependent colocalisation analysis was performed at the level of individual RNP granules, rather than across the whole image. To improve reproducibility, a background-derived thresholding approach was applied for each fluorescence channel. Mean background intensity was quantified from three regions of interest lacking obvious granule signal, and a channel-specific threshold was calculated as follows:

Threshold = mean background intensity + 5 × standard deviation of background intensity

This provided a file-specific background-derived threshold for each channel, rather than relying on manual adjustment of thresholds across images. Thresholded colocalisation analysis was then carried out within individual RNP granule masks. Only RNP granules that remained in focus throughout the movies were selected for further analysis.

For each granule and frame, the following measurements were extracted: RNP granule area, SNRNP70 sum intensity, Manders A colocalisation and Pearson’s coefficient. SNRNP70 sum intensity was additionally normalised to RNP granule area to account for differences in granule size. Where required for comparison across independent experiments, intensity values were min–max normalised to a 0–1 scale prior to plotting and statistical analysis.

### Quantifications and statistical analysis

The number of embryos and samples (n) and definition of statistical significance are indicated in the figure legends. For multiple comparisons, one-way or two-way ANOVA tests with a Sidak correction were used. For RNP granules, statistical analyses were conducted in RStudio using linear mixed-effects models to account for repeated measurements from the same granules over the imaging time course. For time-course analyses of Pearson’s coefficient and colocalisation/Manders A, treatment, frame number and the treatment-by-frame interaction were included as fixed effects, with granule identity included as a random intercept. Models were fitted using maximum likelihood with the lme4 and lmerTest packages, and significance of fixed effects was assessed using Type III ANOVA with Satterthwaite’s approximation for degrees of freedom. The criterion for statistical significance was set at *p* < 0.05 and results are represented as mean ± SEM. Mean values were calculated and plotted using GraphPad Prism.

## Results

### SNRNP70 colocalises with Tardbp in axons

Our prior research demonstrated that the transgenic misexpression of either wild-type or cytosol-restricted human SNRNP70 protein successfully restored motor connectivity and neuromuscular junctions (NMJs) in *snrnp70* null zebrafish (Nikolaou et al., 2022). This functional restoration indicates that human SNRNP70 can interact effectively with the endogenous components of the mRNA processing machinery, including Tardbp. Therefore, we first sought to investigate whether the subcellular distribution of Tardbp overlaps with SNRNP70 in zebrafish. We used an antibody that recognises a conserved region within the C-terminal domain (CTD) of Tardbp protein (Fig.1A). Nucleocytoplasmic fractionation revealed that Tardbp is found not only in its expected location within the nucleus but is also abundant in the cytoplasm (Fig.1B).

**Figure 1:**
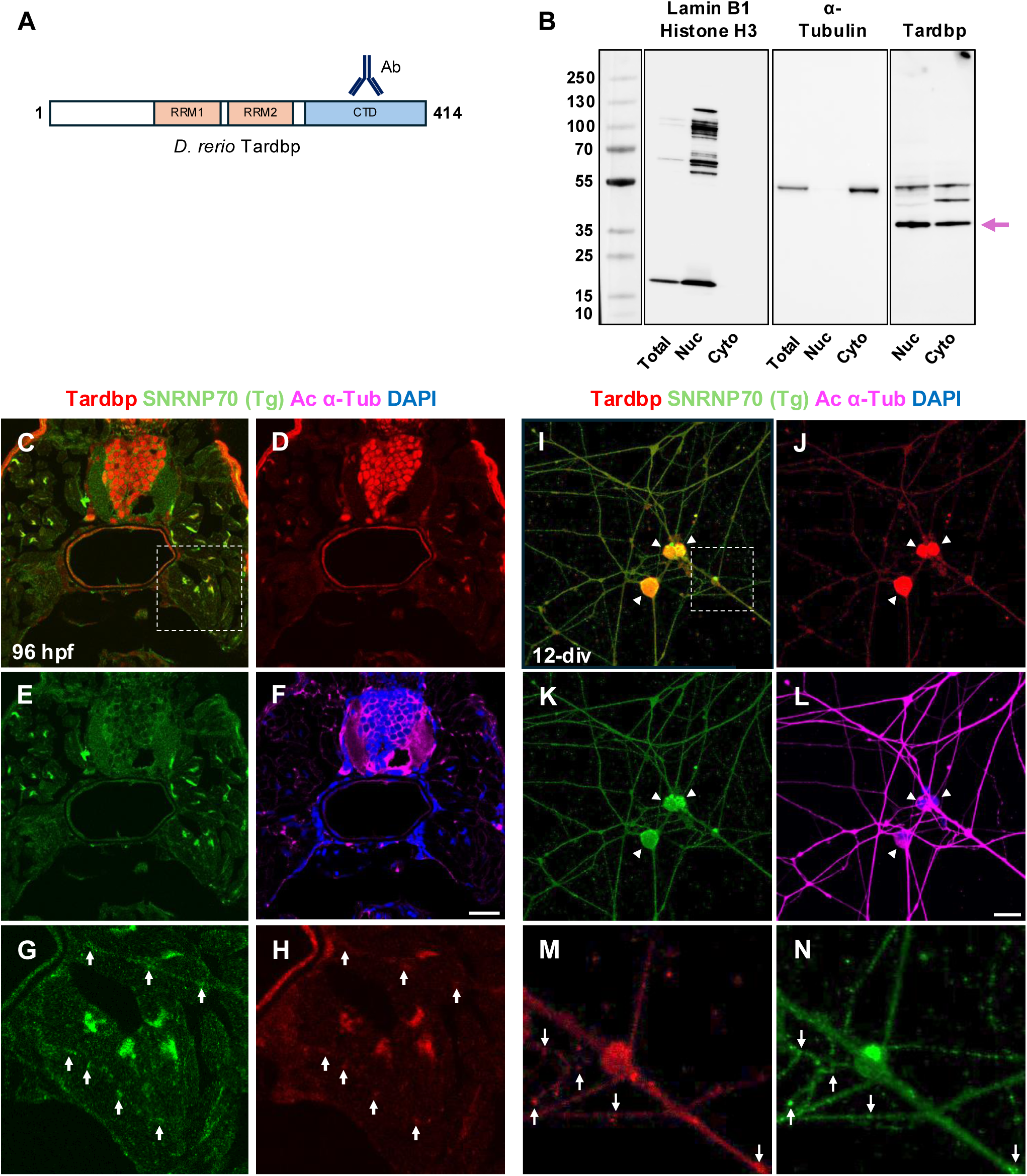
SNRNP70 colocalises with Tardbp in axons. **(A)** Schematic diagram depicting the *D.rerio* Tardbp protein domains: RRM1 (RNA Recognition Motif 1); RRM2 (RNA Recognition Motif 2); and CTD (C-terminus Domain). The location of the primary antibody used is also indicated. **(B)** Immunoblot analysis of Tardbp expression in total, nuclear, and cytoplasmic compartments following subcellular fractionation. Tardbp protein is indicated with a magenta arrow at 43 kDa. A Lamin B1/Histone H3 cocktail was used as a nuclear marker (Histone H3 is predicted to be around 15kDa and LaminB1 detects multiple isoforms between 60-120 kDa), whereas α-Tubulin is a cytoplasmic protein. Protein lysates were produced from 96 hpf wild-type larvae. **(C-H)** Immunohistochemistry on cryosections produced from 96 hpf *Tg(ubi:ERT2:Gal4);(UAS:hSNRNP70:eGFP)* larvae. Antibodies against GFP (to detect the transgenic human SNRNP70 protein), Tardbp and acetylated α-Tubulin were used as indicated together with DAPI for nuclear counterstain. The inset in (C) depicts the regions shown in panels (G) and (H). Arrows indicate peripheral axonal regions where the two proteins appear to colocalise. Scale bar, 20 μm. **(I-N)** Immunocytochemistry on zebrafish embryonic primary neurons at 12 days *in vitro* derived from *Tg(ubi:ERT2:Gal4);(UAS:hSNRNP70:eGFP)* larvae. Antibodies against GFP (to detect the transgenic hSNRNP70 protein), Tardbp and acetylated α-Tubulin were used as indicated together with DAPI for nuclear counterstain. The inset in (I) depicts the regions shown in panels (M) and (N). Arrowheads mark the soma. Arrows indicate axonal puncta where the two proteins appear to colocalise. Scale bar, 10 μm.

We next sought to determine the spatial distribution of Tardbp in neurons, focusing first on the developing spinal cord and peripheral motor axons at 96 hpf. To do this, we utilised a 4-hydroxytamoxifen (4-OHT) inducible transgenic model misexpressing wild-type human SNRNP70 fused to eGFP (Nikolaou et al., 2022), thereafter called SNRNP70 (Tg). Acetylated α-Tubulin (Ac α-Tub) was used to identify neuronal soma and their axons. As anticipated, immunostaining of cryosections revealed a strong nuclear localisation of Tardbp within the spinal cord (Fig.1C-1F). Interestingly, Tardbp was found in distinct axonal puncta colocalising with SNRNP70 (Tg) in peripheral axons (arrows in Fig.1G and 1H).

To characterise the subcellular localisation of Tardbp in more mature neurons, zebrafish embryonic primary neurons derived from the SNRNP70 (Tg) model were utilised following culture for 12 days in vitro (div). Tardbp was prominently detectable in the nucleus and soma cytoplasm (arrowheads in Fig.1I-L). A diffuse and occasionally punctate distribution of Tardbp was also clear within axons (Fig.1J), which appeared to colocalise with SNRNP70 (Tg) (arrows in Fig.1M and 1N). These results indicate that SNRNP70 and TDP-43 colocalise within axonal projections.

### SNRNP70 (Tg) co-associates with Tardbp

To determine whether SNRNP70 interacts with Tardbp, we performed protein co-immunoprecipitation (co-IP) experiments. Using our transgenic misexpression system, we were able to detect two distinct bands between 70-100 kDa in both nuclear and cytoplasmic fractions, corresponding to the SNRNP70 (Tg) protein (arrows in SFig.1A). As a control, a *Tg(huC:GFP*) reporter line was used (in this line GFP is not fused to a protein), confirming the absence of these two bands (SFig.2B). To assess the association between SNRNP70 (Tg) and Tardbp proteins, a total lysate containing the misexpressed SNRNP70 (Tg) protein was used. SNRNP70 (Tg) was immunoprecipitated and detected into the eluate as two clear bands between 70-100 kDa (arrows in Fig.2A). Interestingly, Tardbp was co-immunoprecipitated with SNRNP70, denoted by a single band corresponding to the expected size of the protein at 43 kDa (arrow in Fig.2B). In contrast, an an IgG negative control did not immunoprecipitate Tardbp (Fig.2A and 2B). The findings here indicate that Tardbp interacts with SNRNP70 in our transgenic misexpression system.

**Figure 2:**
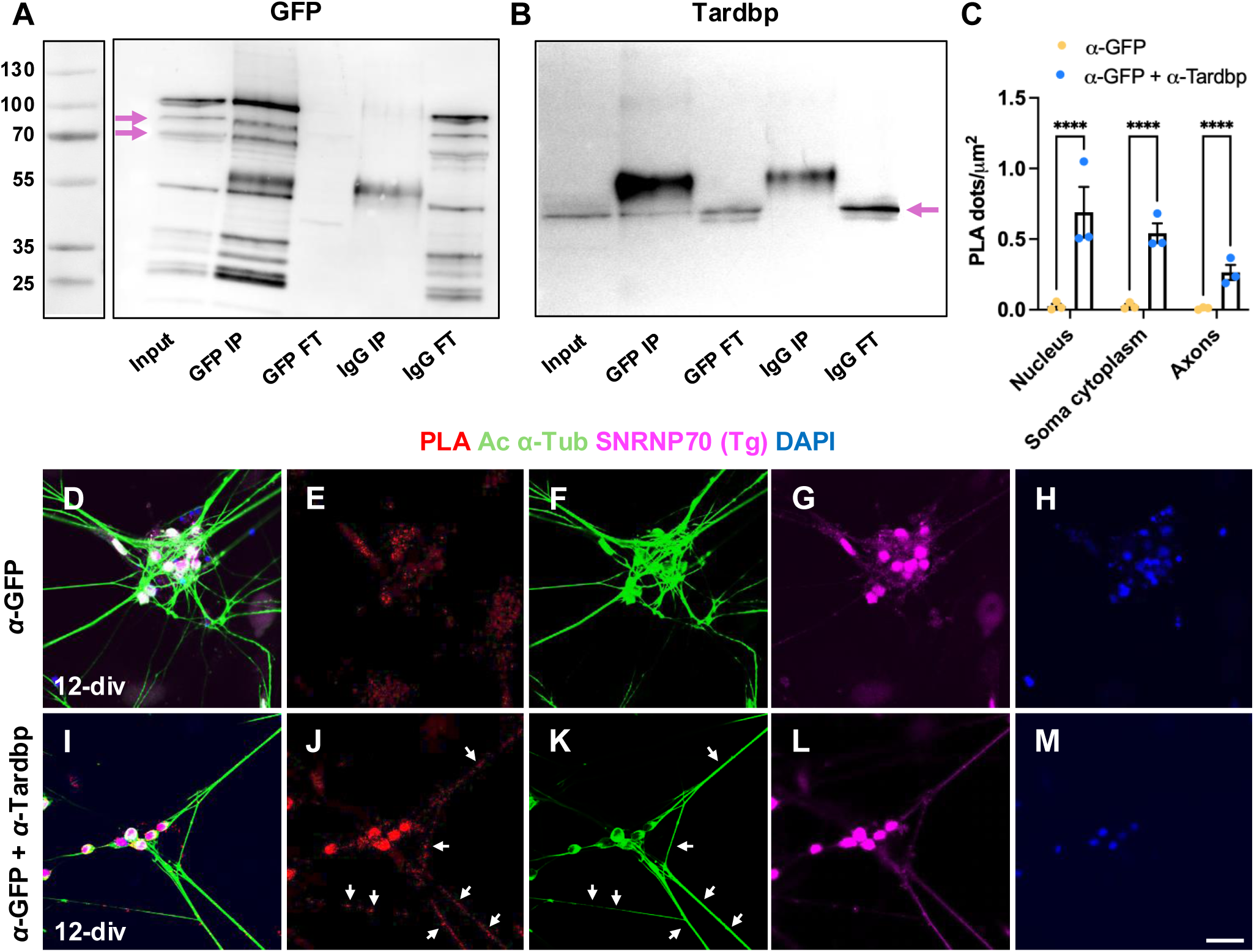
SNRNP70 (Tg) co-associates with Tardbp. **(A)** An immunoblot staining against GPF to detect the presence of transgenic human SNRNP70 protein in an input derived from *Tg(ubi:ERT2:Gal4);(UAS:hSNRNP70:eGFP)* larvae at 4 dpf as well as the immunoprecipitation experiments as labelled. Magenta arrows mark the bands corresponding to SNRNP70 (Tg) protein (between 70-100 kDa). IP: immunoprecipitation; FT: flowthrough. **(B)** An immunoblot staining against Tardbp to detect the presence of this protein in an input derived from *Tg(ubi:ERT2:Gal4);(UAS:hSNRNP70:eGFP)* larvae at 4 dpf as well as the immunoprecipitation experiments as labelled. Magenta arrow marks the band corresponding to Tardbp protein (43 kDa). IP: immunoprecipitation; FT: flowthrough. **(C)** Quantification of the PLA counts using either a GFP antibody alone as a control or both GFP and Tardbp antibodies to detect interactions. PLA was performed on embryonic primary neurons at 12 days *in vitro* derived from *Tg(ubi:ERT2:Gal4);(UAS:hSNRNP70:eGFP)* larvae. PLA dots were counted in the nucleus, soma cytoplasm, and axons before normalising these to the total surface area of the indicated cellular compartment. The graph shows mean values ± SEM. \*\*\*\**p* < 0.0001, two-way ANOVA test with a Sidak correction for multiple comparisons, n = 3 regions of interest per group in three independent experiments. **(D-M)** Representative images showing the distribution of PLA dots in zebrafish embryonic primary neurons at 12 days *in vitro* derived from *Tg(ubi:ERT2:Gal4);(UAS:hSNRNP70:eGFP)* larvae. PLA dots using either a GFP antibody alone as a control (D-H) or both GFP and Tardbp antibodies to detect interactions (I-M) are shown. Antibodies against GFP (to detect the transgenic human SNRNP70 protein) and acetylated α-Tubulin were used as indicated together with DAPI for nuclear counterstain. Arrows in (J) and (K) point towards PLA dots localising to axonal regions. Scale bar, 20 μm.

To map the location of these protein-protein interactions within cells, we performed proximity ligation assays (PLA) on 12-div zebrafish embryonic primary neurons derived from the SNRNP70 (Tg) line. Primary neurons were chosen for efficient image segmentation into subcellular regions (nucleus, soma cytoplasm and axons) and robust quantification of PLA signal. The pipeline of analysis used to identify and count PLA dots within subcellular compartments is outlined in SFig.2. PLA enabled the detection and amplification of fluorescent signals at specific loci within cells reporting the interaction (<40 nm) between Tardbp and GFP (fused to SNRNP70).

An anti-GFP control PLA experiment, involving the use of mock-treated primary neurons without incubation with 4-OHT, revealed a scattering of PLA dots, mainly in the soma (SFig.2C-2G). To note, there was a small amount of SNRNP70 (Tg) protein, as seen in SFig.2F, which is likely to be due to leakiness of the transgenic system. Similarly, an anti-GFP PLA control in 4-OHT-treated primary neurons revealed a salt a pepper pattern of PLA signals, mainly in the soma (Fig.2D-2H, SFig.2H-2L). Therefore, an anti-GFP PLA in 4-OHT treated primary neurons was used as a control to quantify the background amount of PLA. Our experimental PLA group showed an extensive network of SNRNP70 (Tg)-Tardbp interactions across all neuronal compartments, with the highest number of PLA dots being concentrated in the nucleus and soma cytoplasm (Fig.2I-M). Quantifications showed that SNRNP70 (Tg)-Tardbp interactions are widespread within primary neurons with a significant number of interactions happening in the nucleus and soma cytoplasm (Fig.2C). Intriguingly, many interactions localised within axonal projections (arrows in Fig.2J and 2K). PLA dots appeared to coincide with SNRNP70 (Tg) protein distribution in axons (arrowheads in SFig.2O and 2P), further supporting the validity of our PLA results. In conclusion, our observations here support the idea that SNRNP70 and Tardbp co-associate within neurons, including the axonal compartment.

### Endogenous pool of SNRNP70 co-associates with Tardbp in axons

Next, we sought to investigate whether Tardbp interacts with the endogenous pool of SNRNP70 protein. To identify and map locations of such protein-protein interactions, we took advantage of a *TgKI(snrnp70-eGFP)* line recently generated by our group (Lloyd-Jones et al; in preparation). This knock-in line was made by inserting the *egfp* sequence into the endogenous *snrnp70* locus, producing an eGFP fusion at the C-terminus of SNRNP70, thus reporting the localisation and dynamics of the endogenous protein. We performed PLA as above on 12-div zebrafish embryonic primary neurons derived from the *TgKI(snrnp70-eGFP)* line. The assay revealed the presence of several PLA dots in our experimental (anti-GFP + anti-Tardbp) group (Fig.3K and 3P), whereas only few PLA dots could be seen in the anti-GFP control group (Fig.3A and 3F). Careful examination of PLA signal indicated the clear presence of PLA dots in the nucleus (cyan arrowheads in Fig.3L and 3M), soma cytoplasm (yellow arrowheads in Fig.3L-3M), and axons (arrows in Fig.3Q and 3R). In contrast, anti-GFP control images revealed a virtual absence of PLA dots from all cellular compartments (Fig.3B-E and 3G-3J). Quantifications revealed that SNRNP70-Tardbp interactions are indeed significant across all neuronal compartments (Fig.3U). The immunofluorescence data here indicate that Tardbp is not only able to co-associate and interact with a misexpressed transgenic human SNRNP70 protein, but also with the endogenous pool of this protein in zebrafish neurons, including within axons.

**Figure 3:**
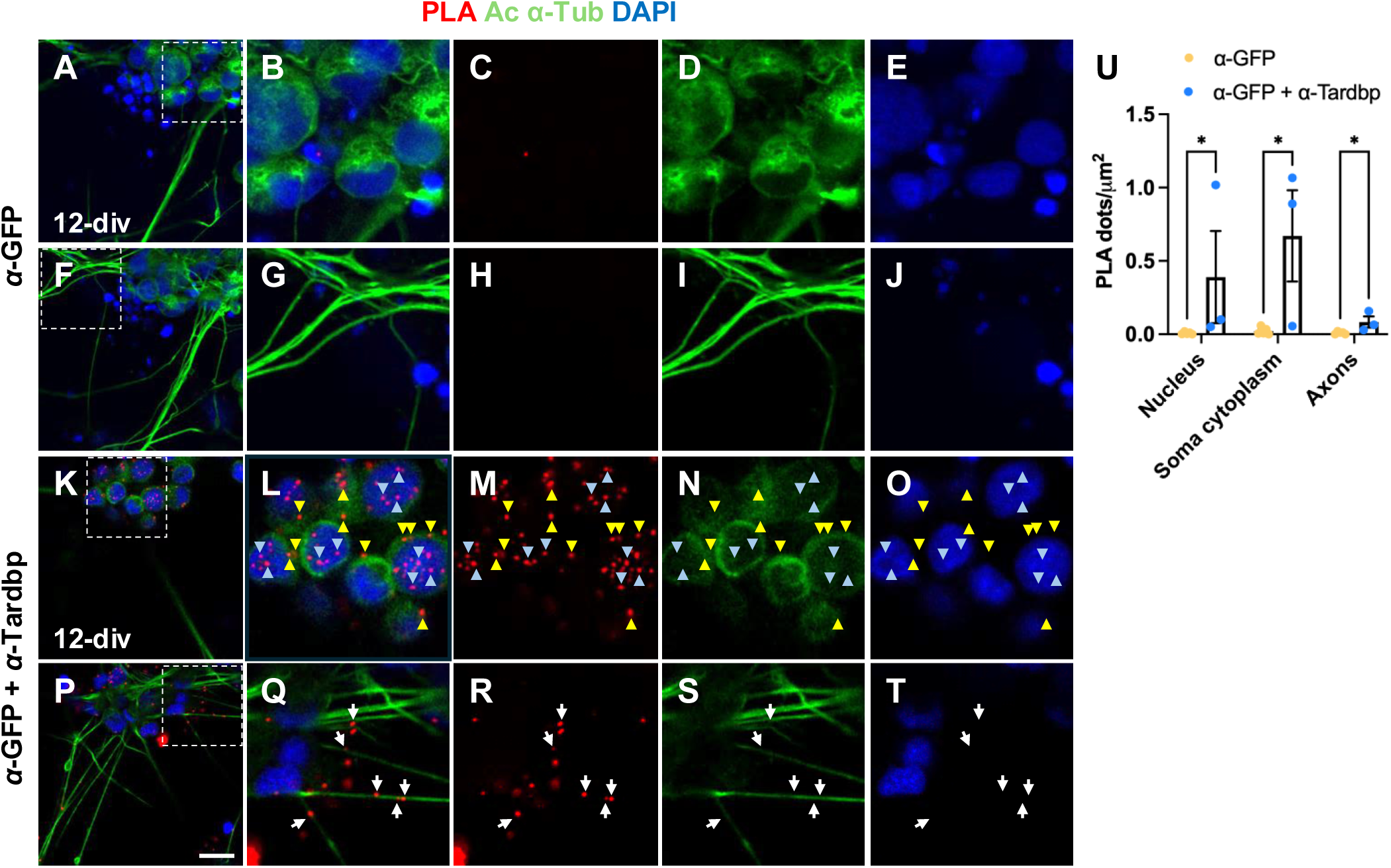
Endogenous pool of SNRNP70 interacts with Tardbp in axons (A-T) Representative images showing the distribution of PLA dots in zebrafish embryonic primary neurons at 12 days *in vitro* derived from *TgKI(snrnp70-eGFP)* larvae. PLA dots using either a GFP antibody alone as a control (A-J) or both GFP and Tardbp antibodies to detect interactions (K-T) are shown. An antibody against acetylated α-Tubulin was used as indicated together with DAPI for nuclear counterstain. The insets shown in (B-E), (G-J), (L-O), and (Q-T) represent the regions marked by dashed squares in panels (A), (F), (K), and (P), respectively. Cyan and yellow arrowheads in panels (L-O) indicate PLA dots localising within the nucleus and soma cytoplasm, respectively. Arrows in panels (Q-T) mark PLA dots found within axons. Scale bar, 10 μm. **(U)** Quantification of the PLA counts using either a GFP antibody alone as a control or both GFP and Tardbp antibodies to detect interactions. PLA was performed on embryonic primary neurons at 12 days *in vitro* derived from *TgKI(snrnp70-eGFP)* larvae. PLA dots were counted in the nucleus, soma cytoplasm, and axons before normalising these to the total surface area of the indicated cellular compartment. The graph shows mean values ± SEM. \**p* < 0.05, two-way ANOVA test with a Sidak correction for multiple comparisons, n = 3 regions of interest per group from three independent experiments.

### *snrnp70* interacts genetically with *tardbp*

We have previously shown that *snrnp70^kg163^* null zebrafish embryos display motor axonal outgrowth and neuromuscular defects, which are partially restored following cytosolic SNRNP70 misexpression (Nikolaou et al., 2022). This observation suggests that although the neuromuscular phenotype is not exclusively triggered by the loss of non-nuclear SNRNP70, this cytoplasmic pool provides an important contribution to the development of neuromuscular connectivity by regulating aspects of motor axonal growth and synaptogenesis. To examine whether SNRNP70 could participate in the regulation of mRNA processing alongside Tardbp, we first sought to investigate whether their loss-of-function results in similar motor axonal defects. To generate *tardbp* F0 knock-outs of, a cocktail of three sgRNAs (SFig.3A) was delivered into fertilised eggs. This approach resulted in a deletion of a large genomic DNA region in the middle of *tardbp* gene (SFig.3B) and a marked reduction in the amount of Tardbp protein produced (SFig.3C), validating the effectiveness of our crispant approach. Phenotypic analysis showed that, on average, *tardbp* crispants had a motor nerve length of 105 μm (Fig.4B), which is significantly shorter than the average length of wild-type (150 μm) control embryos at 30 hpf (Fig.4A and 4E).

**Figure 4:**
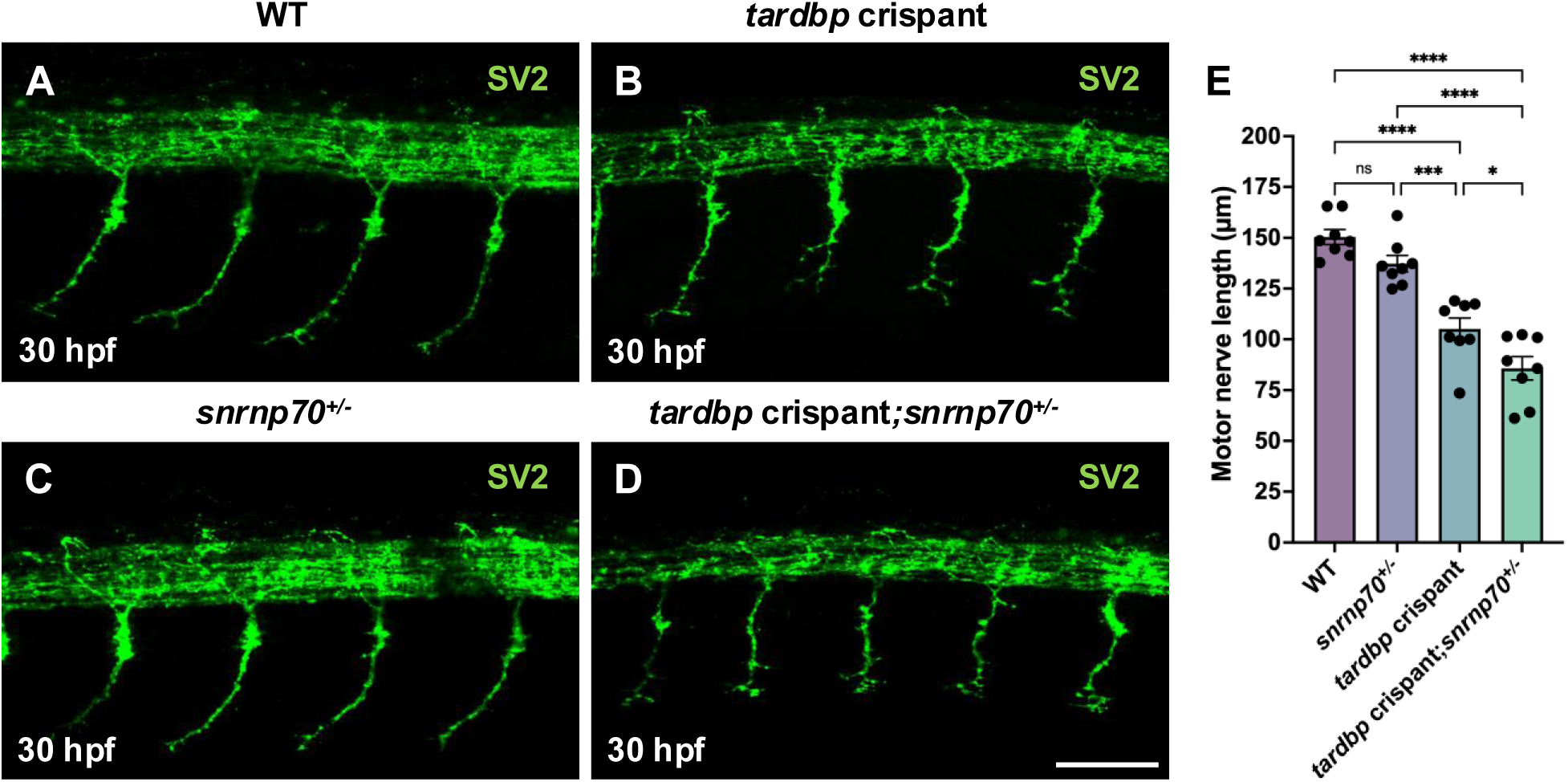
snrnp70 interacts genetically with tardbp (A-D) Representative images showing anti-SV2 immunostaining marking zebrafish motor neuron nerves at 30 hpf in wild-type (A), *tardbp* crispant knock-out (B), heterozygous *snrnp70* (C), and *tardbp* crispant knock-out on a heterozygous *snrnp70* background (D). Scale bar, 50 μm. **(E)** Quantification of the motor neuron nerve length in the four experimental groups as indicated. The graph shows mean values ± SEM. \*\*\*\**p* < 0.0001; \*\*\**p* < 0.001; \**p* < 0.05; ns, not significant, one-way ANOVA test with a Sidak correction for multiple comparisons, n = 8 animals per group from two independent experiments.

The similarity of the motor axonal growth defects following *snrnp70* and *tardbp* loss-of-function, raised the possibility that SNRNP70 may interact with Tardbp. We therefore examined whether *snrnp70* and *tardbp* interact genetically. Heterozygous *snrnp70^kg163^* carriers had normal motor axon extensions, with an average motor nerve length of 137 μm (Fig.4C). The average motor nerve length was significantly shorter in larvae heterozygous for *snrnp70^kg163^* that also lacked *tardbp* (Fig.4D). Here, the average length of 85 μm was significantly lower than *snrnp70^kg163^* We therefore sought to determine whether the recruitment of SNRNP70 to these RNP granules is dependent on Tardbp. TDP-43 is a well-characterised component of RNP transport granules in neurons that binds to specific target mRNAs and facilitates their incorporation into such membrane-less organelles (Alami et al., 2014). the two proteins contribute to the same biological pathways and are involved in common processes essential for motor neuron development.

### Tardbp facilitates the recruitment of SNRNP70 to RNP granules

Previous findings indicated that transgenic SNRNP70 protein localises to RNP granules in zebrafish neurons (Nikolaou et al., 2022). We therefore sought to determine whether the recruitment of SNRNP70 to these RNP granules is dependent on Tardbp. TDP-43 is a well-characterised component of RNP transport granules in mammalian neurons that binds to specific target mRNAs and facilitates their incorporation into such membrane-less organelles (Alami et al., 2014).

To monitor the localisation of endogenous SNRNP70 within RNP granules, we used our *TgKI(snrnp70-eGFP)* line. To visualize RNP granules, we used a fluorescent derivative of uridine-5’-triphosphate (Cy5-UTP) that is incorporated into RNA molecules. The Cy5-labelled RNA is eventually incorporated into cytoplasmic RNP granules, thus labelling these with Cy5 (Nikolaou et al., 2022; Wong et al., 2017). We observed several cytoplasmic eGFP^+^ puncta within spinal cord neurons at 52 hpf, which report the distribution of SNRNP70, frequently colocalising with Cy5-RNA^+^ RNP granules outside the neuronal nucleus. The majority of the eGFP^+^ puncta appeared motile (as shown in Video S1 and S2), indicating that SNRNP70 is incorporated into motile RNP granules.

To determine whether the ability of SNRNP70 to localise to these RNP granules is dependent on Tardbp, we generated *tardbp* F0 knock-outs in the *TgKI(snrnp70-eGFP)*/Cy5-RNA^+^ RNP background. We first examined whether loss of *tardbp* alters the characteristics of RNP granules and found that their size was significantly larger in the crispant than to the wild-type group (Fig.5M). These results suggest that Tardbp may regulate aspects of RNP granule biogenesis. Analysis of SNRNP70 fluorescence within these granules revealed that the amount of SNRNP70 inside RNP granules is lower in the *tardbp* crispants, however, pairwise frame-wise comparisons did not remain significant after correction for multiple comparisons (Fig.5N).

**Figure 5:**
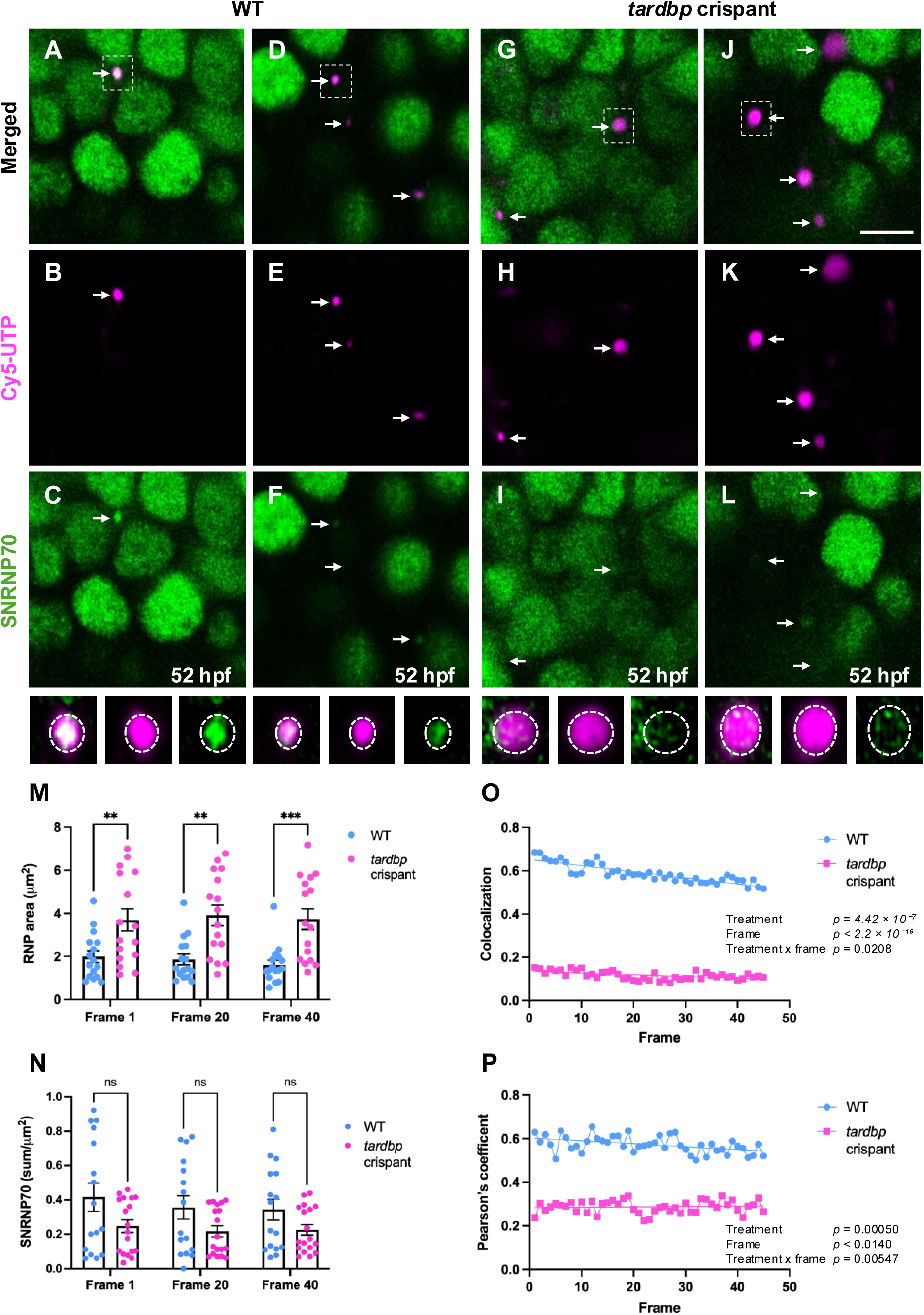
Tardbp is required for SNRNP70 recruitment to RNP granules (A-L) Representative images (all at frame 1) taken from short time-lapse movies of spinal cord neurons in the *TgKI(snrnp70-eGFP)* line at 52 hpf. Cy5-UTP was used as a marker of RNP granules. Arrows indicate RNP granules in the spinal cord of wild-type (A-F) and *tardbp* crispant knock-out (G-L) animals. Bottom: insets representing the regions marked by a dashed squares in panels (A), (D), (G), and (J). Scale bar, 5 μm. **(M)** Quantification of the RNP granule size at the start (frame 1), middle (frame 20) and towards the end (frame 40) of the time-lapse imaging in wild-type and *tardbp* crispant knock-outs. The graph shows mean values ± SEM. \*\*\**p* < 0.001; \*\**p* < 0.01, two-way ANOVA test with a Sidak correction for multiple comparisons, n = 16 RNP granules per group, wild-type (n = 4) and *tardbp* crispant knock-out (n = 5) animals from two independent experiments. **(N)** Quantification of the SNRNP70-eGFP fluorescence (sum values) normalised to RNP granule size at the start (frame 1), middle (frame 20) and towards the end (frame 40) of the time-lapse imaging in wild-type and *tardbp* crispant knock-outs. The graph shows mean values ± SEM. ns, not significant, two-way ANOVA test with a Sidak correction for multiple comparisons, n = 16 RNP granules per group, wild-type (n = 4) and *tardbp* crispant knock-out (n = 5) animals from two independent experiments. **(O)** Quantification of SNRNP70-eGFP and Cy5-RNA granule colocalisation over time in wild-type and *tardbp* crispant knock-out granules. Colocalisation was measured within masked RNP granule regions across the time-lapse series. Linear regression lines show the fitted temporal trend for each genotype. Linear mixed-effects modelling showed that colocalisation was significantly lower in *tardbp* crispant knock-out granules than in wild-type across the imaging time course, and that the temporal profile of colocalisation differed between the two groups. Data are from 16 RNP granules per group, derived from wild-type (*n* = 4) and *tardbp* crispant knock-out (*n* = 5) animals across two independent experiments. Treatment effect: *p* = 4.42 × 10⁻⁷; frame effect: *p* < 2.2 × 10⁻¹⁶; treatment × frame interaction: *p* = 0.0208. **(P)** Quantification of the linear relationship between SNRNP70-eGFP and Cy5-RNA granule fluorescence over time in wild-type and *tardbp* crispant knock-out granules. Pearson’s coefficient was measured within masked RNP granule regions across the time-lapse series. Linear regression lines show the fitted temporal trend for each genotype. Linear mixed-effects modelling showed that Pearson’s coefficient was significantly lower in *tardbp* crispant knock-out granules than in wild-type across the imaging time course, and that the temporal profile differed between the two groups. Data are from 16 RNP granules per group, derived from wild-type (n = 4) and *tardbp* crispant knock-out (n = 5) animals across two independent experiments. Treatment effect: *p* = 0.00050; frame effect: *p* = 0.0140; treatment × frame interaction: *p* = 0.00547.

We wondered whether the spatial distribution of SNRNP70 within RNP granules is altered following the depletion of Tardbp. Analysis of individual RNP granules showed sustained moderate-to-high colocalisation between the Cy5-RNA and SNRNP70 signals in the wild-type group over time (SFig.4A), with an estimated Manders A value of 0.58 ± 0.05 at the centred frame position (Fig. 5O). By contrast, colocalisation was markedly diminished in *tardbp* crispant granules (SFig.4B), with an estimated Manders A value of 0.13 ± 0.05 (Fig.5O). This difference in colocalisation between SNRNP70 and Cy5-RNA was sustained across time, between the *tardbp* crispant relative to the wild-type group (p = 4.42 × 10⁻⁷) (Fig.5O). The behaviour of colocalisation was also significantly different across frame numbers (p < 2.2 × 10⁻¹⁶), with a significant treatment-by-frame interaction (p = 0.0208) (Fig. 5O), indicating that the temporal profile of colocalisation differed between wild-type and crispant groups. Examination of the relationship between SNRNP70-eGFP and Cy5-RNA signal intensities (Pearson’s coefficient) of individual RNP granules revealed stronger positive linear relationships over time between the two fluorescent signals in the wild-type compared to crispant group (SFig.5A and 5B). Pearson’s coefficient was significantly reduced in *tardbp* crispant granules compared to wild-type granules across the imaging time course (*p* = 0.00050) (Fig.5P). A significant treatment-by-frame interaction was also detected (*p* = 0.00547) (Fig.5P), indicating that the temporal profile of Pearson’s coefficient differed between the two groups. Taken together, these findings suggest that Tardbp is required for the recruitment and localisation of SNRNP70 to RNP granules.

### SNRNP70 and TDP-43 interact in human iPSC-derived motor neurons

Our current findings show an intriguing co-association between Tardbp and SNRNP70, with Tardbp appearing to be essential for the recruitment of SNRNP70 to RNP granules. Given that RBPs have conserved roles in regulating the transport and processing of mRNAs within such membrane-less organelles, we hypothesised that the interactions between Tardbp and SNRNP70 are not unique to zebrafish neurons but evolutionarily conserved across species. To address this, we used matured human induced pluripotent stem cell (hiPSCs) derived motor neurons, which have endogenously tagged TDP-43 with a C-terminal GFP (Ganssauge et al., 2025), and performed PLA to identify and map interactions between human TDP-43 and SNRNP70 proteins. For the experimental group, antibodies against GFP and SNRNP70 were used, showing sparse PLA dots in the soma (Fig.6B). Intriguingly, PLA dots can also be seen within Neurofilament-positive axons (arrows in Fig.6C and 6D). In contrast, only few PLA dots could be seen in the anti-GFP control group (Fig.6A). Our quantification revealed that the number of PLA dots was significantly higher in all neuronal compartments compared to the anti-GFP control group (Fig.6E). Collectively, these findings indicate that neuronal SNRNP70-TDP-43 co-associations are evolutionary conserved, raising the possibility that these two proteins may have a conserved function in neurons generally and RNP granules more specifically.

**Figure 6:**
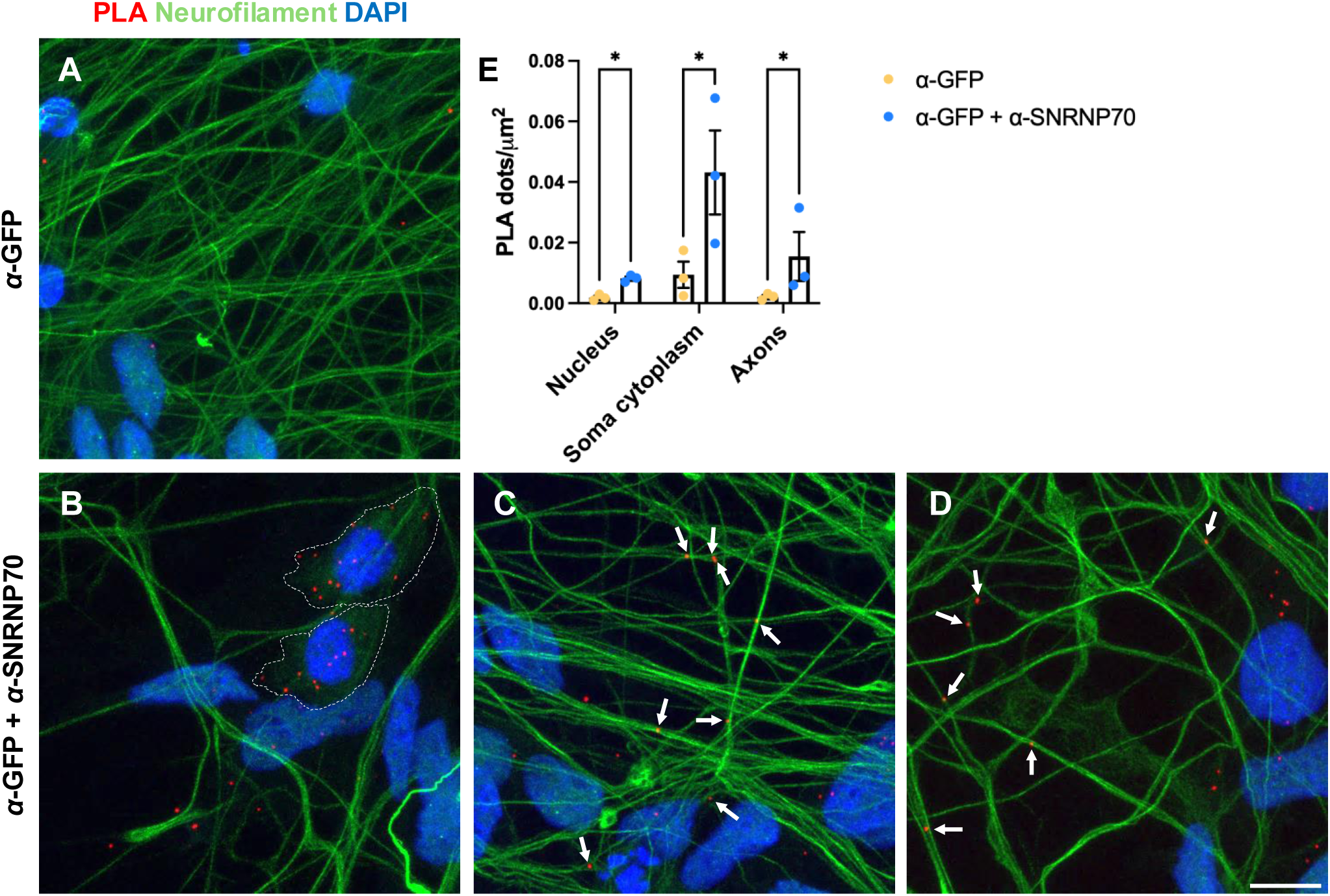
SNRNP70 and TDP-43 interact in human iPSC-derived motor neurons (A-D) Representative images showing the distribution of PLA dots in mature human motor neurons derived from engineered TDP43-GFP iPSCs. PLA dots using either a GFP antibody alone as a control (A) or both GFP and SNRNP70 antibodies to detect interactions (B-D) are shown. An antibody against Neurofilament was also used together with DAPI for nuclear counterstain. The dashed area in panel (B) marks the soma region of the labelled cells. Arrows in panels (C) and (D) mark PLA dots found within Neurofilament-positive axonal regions. Scale bar, 10 μm. **(E)** Quantification of the PLA counts using either a GFP antibody alone as a control or both GFP and SNRNP70 antibodies to detect interactions. PLA was performed on mature human motor neurons derived from engineered TDP43-GFP iPSCs. PLA dots were measured in the nucleus, soma cytoplasm, and axons before normalising these to the total surface area of the indicated cellular compartment. The graph shows mean values ± SEM. \**p* < 0.05, two-way ANOVA test with a Sidak correction for multiple comparisons, n = 3 regions of interest per group from three independent experiments.

## Discussion

Previous studies have identified protein interactions between TDP-43 and SNRNP70 in the context of neurodegeneration (Bishof et al., 2018; Ling et al., 2010) and aging (Rhine et al., 2025) using biochemical approaches. Our work here shows for the first time the existence of associations between these two proteins under physiological conditions during development of neuronal connections. Whilst a large number of interactions are found within the neuronal nucleus, supporting a shared role in pre-mRNA splicing (Brown et al., 2022; Huranová et al., 2010), many associations are also mapped to the soma cytoplasm and axonal projections. The intriguing diversity of extra-nuclear interactions suggests a shared role in mRNA processing beyond what is classically associated with RNA splicing regulators. Indeed, our work here proposes a model where SNRNP70, likely through direct interactions with TDP-43, is recruited to RNP granules to regulate the extra-nuclear processing of mRNA transcripts.

Our findings point to a physiological role of TDP-43 protein interactions with SNRNP70 and a shared role during neuronal development. TDP-43 acts as a master regulator of mRNA processing. Beyond its well-established role in pre-mRNA splicing, TDP-43 has a pivotal role in transporting RNP granules in dendrites, inhibiting translation inside those granules, and reactivating it once the granules reach the dendritic spines (Chu et al., 2019; Majumder et al., 2016). During motor neuron development, TDP-43 is primarily responsible for ensuring the stability and transport of crucial mRNAs, particularly those required for building axons and maintaining synaptic connections (Briese et al., 2020). Similarly, SNRNP70’s main role inside the nucleus as part of the spliceosome has been characterised (Kondo et al., 2015). Intriguingly, SNRNP70 is also required outside the nucleus for the stability and transport of mRNA transcripts in neurons (Nikolaou et al., 2022) and for the apical localisation of transcripts in the gut epithelium (Lee et al., 2025). The shared function of SNRNP70 and TDP-43 is further supported by our findings showing co-associations across several neuronal compartments where mRNA processing takes place. Our genetic interaction results further support the idea that these two proteins act synergistically during motor neuron development.

Interestingly, previous results have shown that loss of Tardbp in zebrafish does not affect spinal motor neuron axon outgrowth because of alternative splicing of a second *TARDBP* ortholog gene called *tardbpl (Hewamaddumal et al., 2013; Schmid et al., 2013)*. The discrepancy between our results and previous findings could be rooted in the gene editing approach used here. In our study, a genomic deletion affecting a large part of the protein was achieved. Nevertheless, loss of both ortholog proteins resulted in motor axon defects (Hewamaddumal et al., 2013; Schmid et al., 2013) that mimic our *tardbp* crispant findings.

Dysregulation of both TDP-43 and SNRNP70 is closely associated with neurodegenerative diseases like Amyotrophic Lateral Sclerosis (ALS), Frontotemporal Lobar Degeneration (FTLD), and Alzheimer’s Disease (AD). In over 97% of ALS cases and roughly half of FTD cases, TDP-43 is mislocalised from the nucleus to the cytoplasm, where it forms toxic, insoluble aggregates (Suk & Rousseaux, 2020). As TDP-43 leaves the nucleus, the cell loses its ability to accurately splice pre-mRNA. This leads to the abnormal retention of cryptic exons in crucial transcripts (such as *UNC13A* and *STMN2*), which destabilises the proteins necessary for synaptic maintenance and axonal growth (Brown et al., 2022; Klim et al., 2019; Ma et al., 2022). The mislocalisation of TDP-43 is also hypothesised to promote G3BP1-positive RNP assembly, which ultimately disrupts global protein synthesis in axons and NMJs and induces mitochondrial dysfunction (Altman et al., 2021). In the case of SNRNP70, its mislocalisation, aggregation, and specific proteolytic cleavage are increasingly recognised as primary drivers of neuronal toxicity, synaptic dysfunction, and axonal degeneration (Bai et al., 2013; Chen et al., 2022). Intriguingly, the two low complexity domains found within the C-terminus of the SNRNP70 protein are necessary for the co-aggregation of SNRNP70 with Tau in both sporadic and familial human cases of AD (Bishof et al., 2018). Moreover, these domains are also essential for interactions with TDP-43 (Bishof et al., 2018). The convergence of TDP-43 and SNRNP70 highlights their essential roles in RNA processing, with defects in these processes serving a central mechanism in neurodegeneration. By disrupting both the nuclear splicing machinery as well as the axonal transport and stability of mRNAs targets, the interplay between these proteins sets off a cascade of neuronal toxicity. Our findings here, mapping the extent of interactions between these two proteins, provide further support for their shared roles in neurodegeneration.

TDP-43 is a key RBP that acts as a scaffold and regulator in the assembly of RNP granules such as stress granules and transport granules. Through liquid-liquid phase separation, it organises target RNAs and other proteins, facilitating dynamic granule assembly, localised mRNA translation, and cellular stress responses (Corbet et al., 2021). Structural motifs that are important for TDP-43 liquid-liquid phase separation include the N-terminal domain enabling self-oligomerisation (Zhang et al., 2013) and the C-terminal domain that is a critical regulator of protein-protein interactions (Buratti et al., 2005; D’Ambrogio et al., 2009). Our findings showing that TDP-43 is required for SNRNP70 localisation to RNP granules further support the idea that TDP-43 acts a master organiser of such membrane-less organelles by participating in the recruitment of key proteins. The phase separation and maintenance of liquid properties also depend on the RNA recognition motifs mediating binding to target mRNAs (Grese et al., 2021). Interestingly, recent *in vivo* findings indicate that RNA-binding deficient TDP-43 variants display reduced cytoplasmic localisation (Scherer et al., 2024), suggesting the formation of cytoplasmic RNP granules depends on mRNA binding. It is likely that by binding to target mRNAs and recruiting SNRNP70 to RNP granules, TDP-43 enables SNRNP70-mediated mRNA processing to take place.

In conclusion, our research reveals a mechanism reliant on TDP-43 that allows SNRNP70 to associate with RNP granules as neuronal connections develop. Future investigations will focus on detailing the specific nature and the comprehensive range of biological roles these protein-protein interactions play in both health and disease.

## Acknowledgements

We thank Dr Julia Sero for assistance in using Cell Profiler. We thank staff at the University of Bath fish facility and University of Exeter Aquarium Research Centre for their fish husbandry and care. We also thank staff at the Bioimaging facility (University of Exeter) for their support.

## Author contributions

Conceptualisation: T.B., N.N. Data curation: T.B., C.M.E., S.M.E.J., J.E., N.N. Formal analysis: T.B., J.L-J., C.M.E., S.M.E.J., J.E., C.L., N.N. Funding acquisition: A.B., N.N. Investigation: T.B., J.L-J., C.M.E., S.M.E.J., J.E., J.G., N.N. Methodology: T.B., J.L-J., C.M.E., S.M.E.J., J.E., C.L., N.N. Project administration: N.N. Resources: T.B., J.L-J., C.M.E., S.M.E.J., J.G. Supervision: T.B., A.B., N.N. Writing – original draft: N.N. Writing – review and editing: T.B., J.L-J., C.M.E., S.M.E.J., J.E., J.G., C.L., A.B., N.N.

---

**Supplementary figure 1: SNRNP70 (Tg) detection on immunoblots**

**(A)** An immunoblot using an antibody against GFP (goat anti-GFP) showing the presence of transgenic human SNRNP70 protein in total, nuclear and cytoplasmic fractions of protein lysates derived from *Tg(ubi:ERT2:Gal4);(UAS:hSNRNP70:eGFP)* larvae at 4 dpf. Magenta arrows mark the bands corresponding to SNRNP70 (Tg) protein (between 70-100 kDa).

**(B)** An immunoblot using an antibody against GFP (rabbit anti-GFP) showing the presence of transgenic human SNRNP70 protein in total, nuclear and cytoplasmic fractions of protein lysates derived from either *Tg(ubi:ERT2:Gal4);(UAS:hSNRNP70:eGFP)* or *Tg(huC:GFP)* larvae at 4 dpf. Magenta arrows mark the bands corresponding to SNRNP70 (Tg) protein (between 70-100 kDa). Note the absence of SNRNP70 (Tg)-specific bands in protein lysates derived from the *Tg(huC:GFP)* reporter line.

**Supplementary figure 2: PLA methodology and optimisation**

**(A)** A representative example from a PLA experiment showing the distribution of PLA signal in zebrafish embryonic primary neurons at 12 days *in vitro* derived from *Tg(ubi:ERT2:Gal4);(UAS:hSNRNP70:eGFP)* larvae. The samples were also counterstained with DAPI (to mark the nucleus) and acetylated α-Tubulin (to mark axons). Scale bar, 20 μm.

**(B)** Pipeline of Cell Profiler analysis that was followed to count PLA dots per neuronal compartment. Scale bar, 20 μm.

**(C-Q)** Representative images showing the distribution of PLA dots in zebrafish embryonic primary neurons at 12 days *in vitro* derived from *Tg(ubi:ERT2:Gal4);(UAS:hSNRNP70:eGFP)* larvae. PLA dots using a GFP antibody alone in mock-treated cells (C-G), GFP antibody alone in 4-OHT-treated cells (H-L), or both GFP and Tardbp antibodies in 4-OHT-treated cells as the experimental group (M-Q) are shown. Antibodies against GFP (to detect the transgenic human SNRNP70 protein) and acetylated α-Tubulin were used as indicated together with DAPI for nuclear counterstain. The insets shown in (D-G), (I-L) and (N-Q) represent the regions marked by a dashed squares in panels (C), (H) and (M), respectively. Arrows in (O) and (P) point towards PLA dots colocalising with SNRNP70 (Tg) protein to axonal regions. Scale bar, 20 μm.

**Supplementary figure 3: Validation of *tardbp* F0 knock-out model**

**(A)** Genomic locus of zebrafish *tardbp*. sgRNA binding sites are indicated. Primers used for genomic amplification in (B) are also shown.

**(B)** Amplification from genomic DNA extracted from uninjected and *tardbp* crispant embryos. Each lane represents amplification from individual embryos.

**(C)** Immunoblots using antibodies against either Tardbp or Histone H3 showing the presence of Tardbp in uninjected, mock-injected and *tardbp* crispant embryos at 48 hpf. Magenta arrow marks the bands corresponding to Tardbp protein (43 kDa).

**Supplementary figure 4: Spatial overlap between Cy5-RNA and SNRNP70 in individual RNP granules**

(A) Representative graphs showing the colocalisation between SNRNP70-eGFP and Cy5-RNA fluorescence over time within individual wild-type RNP granules.

(B) Representative graphs showing the colocalization between SNRNP70-eGFP and Cy5-RNA fluorescence over time within individual *tardbp* crispant knock-out RNP granules.

**Supplementary figure 5: Signal intensity relationship between Cy5-RNA and SNRNP70 in individual RNP granules**

(A) Representative graphs showing the Pearson’s overlap (colocalisation) between SNRNP70-eGFP and Cy5 fluorescence over time within individual wild-type RNP granules.

(B) Representative graphs showing the spatial overlap (colocalisation) between SNRNP70-eGFP and Cy5 fluorescence over time within individual *tardbp* crispant knock-out RNP granules.

**Supplementary video 1: Motile SNRNP70^+^ RNP granule in a wild-type spinal cord**

Representative example of a SNRNP70^+^ RNP granule in the spinal cord of wild-type larva at 52 hpf. The granule appears to be motile during the duration of the video.

**Supplementary video 1: Motile SNRNP70^+^ RNP granule in a *tardbp* KO spinal cord**

Representative example of a SNRNP70^+^ RNP granule in the spinal cord of *tardbp* KO larva at 52 hpf. The granule appears to be motile during the duration of the video.

