## Supplemental figures for "SNRNP70 interacts with TDP-43 to promote RNP granule localisation and regulate motor neuron development"

### Supplementary figure 1: SNRNP70 (Tg) detection on immunoblots

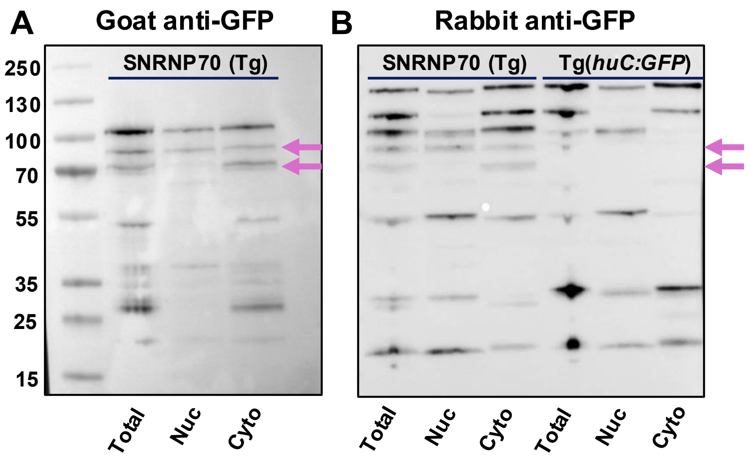

### Supplementary figure 2: PLA methodology and optimisation

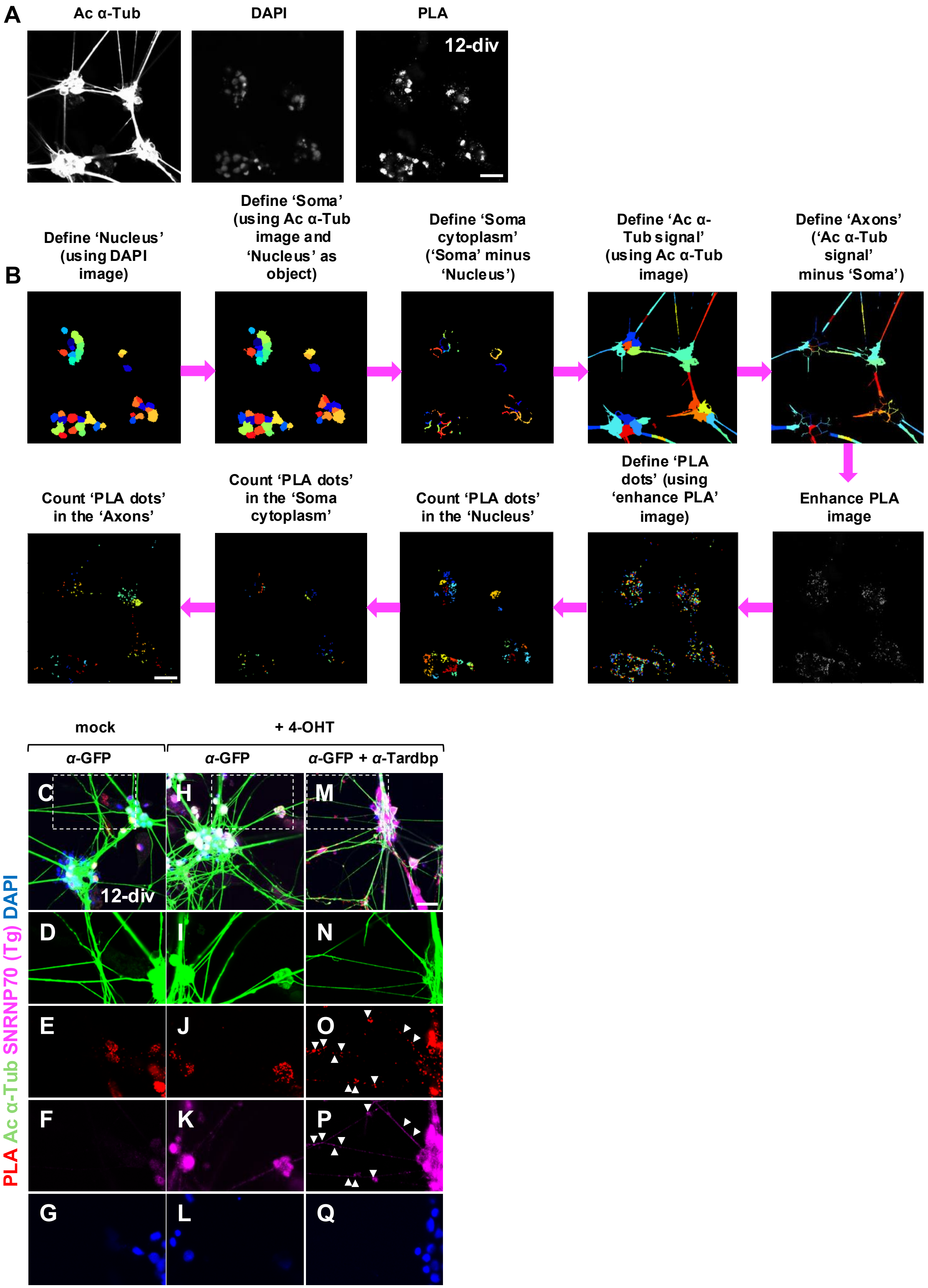

### Supplementary figure 3: Validation of *tardbp* F0 knock-out model

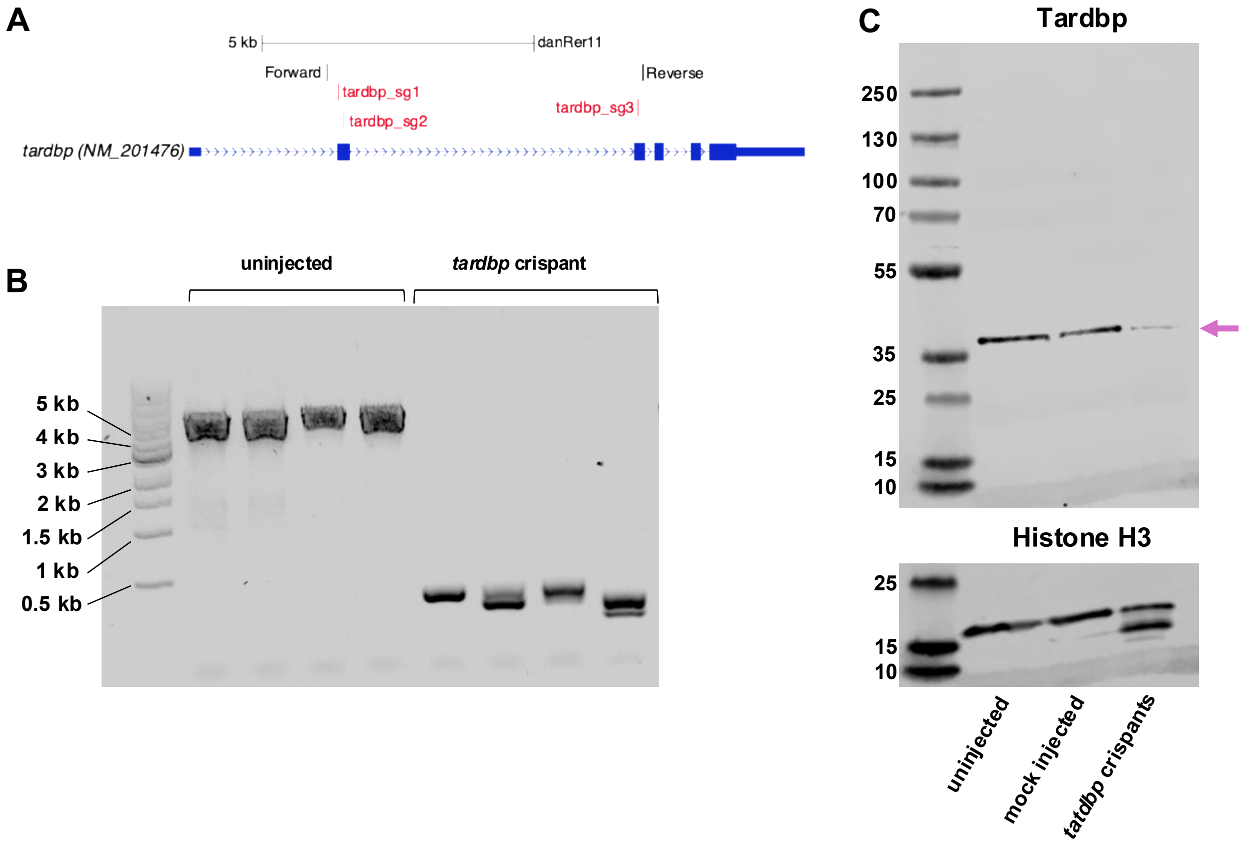

#### Supplementary figure 4: Spatial overlap between Cy5-RNA and SNRNP70 in individual RNP granules

A

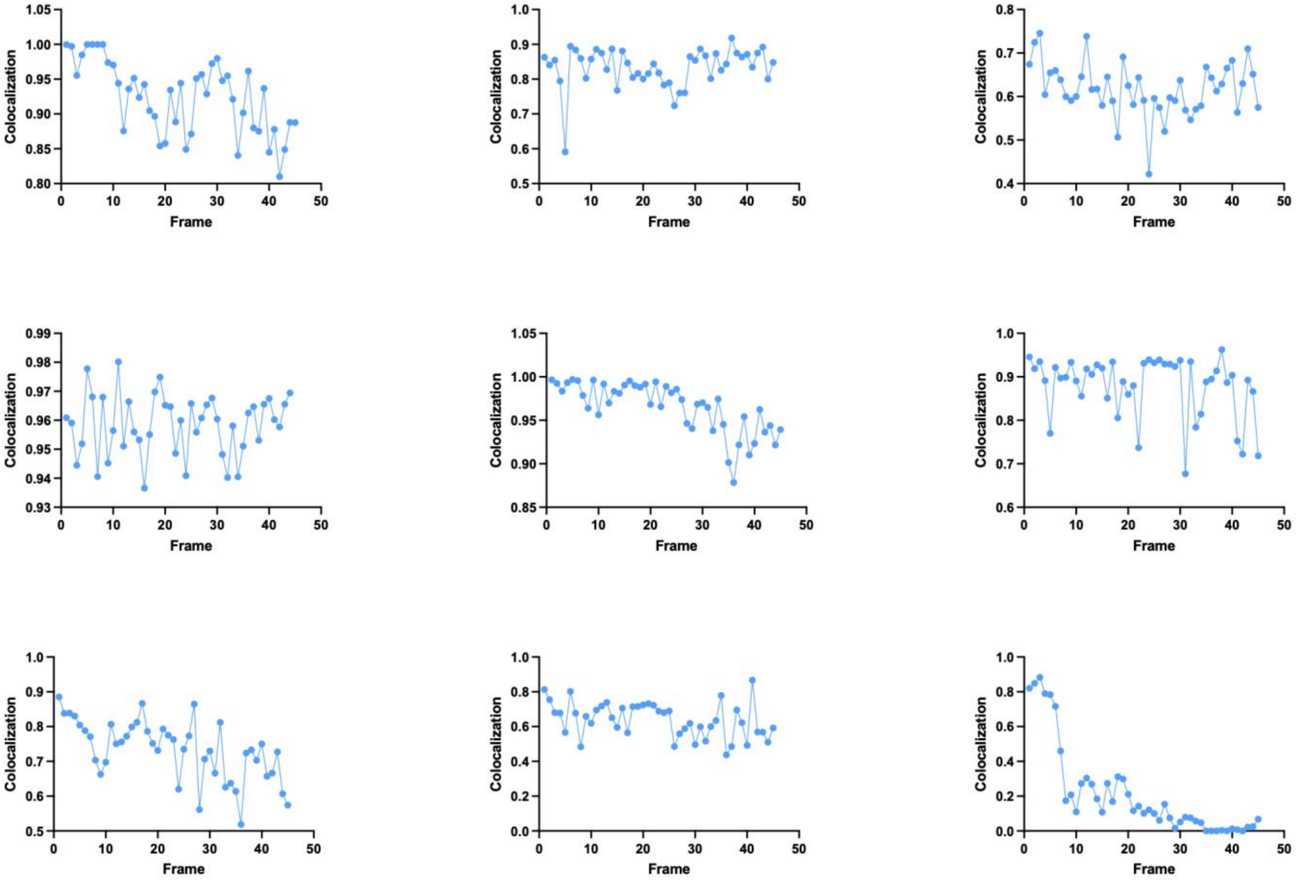


B

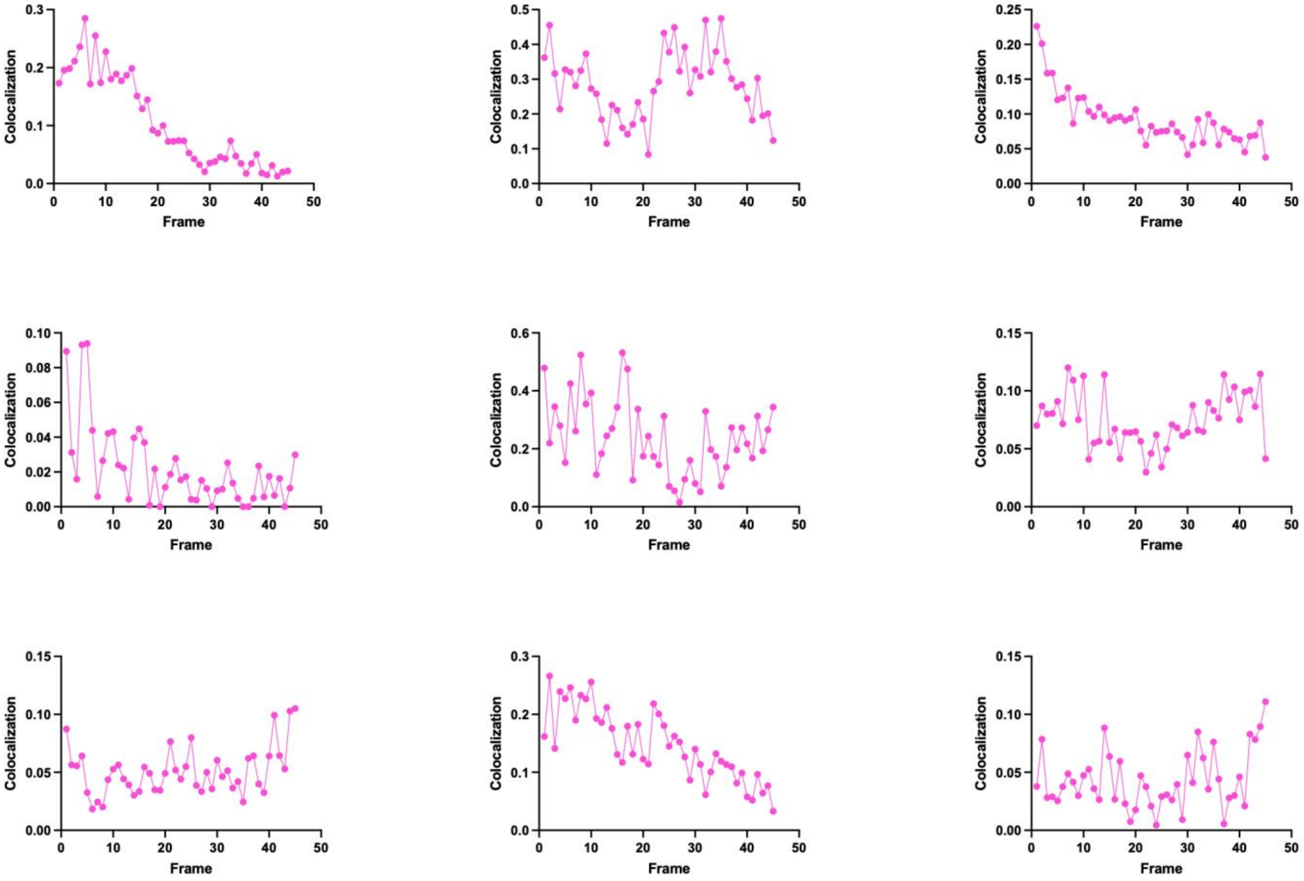

**Supplementary figure 5: Signal intensity relationship between Cy5-RNA and SNRNP70 in individual RNP granules**

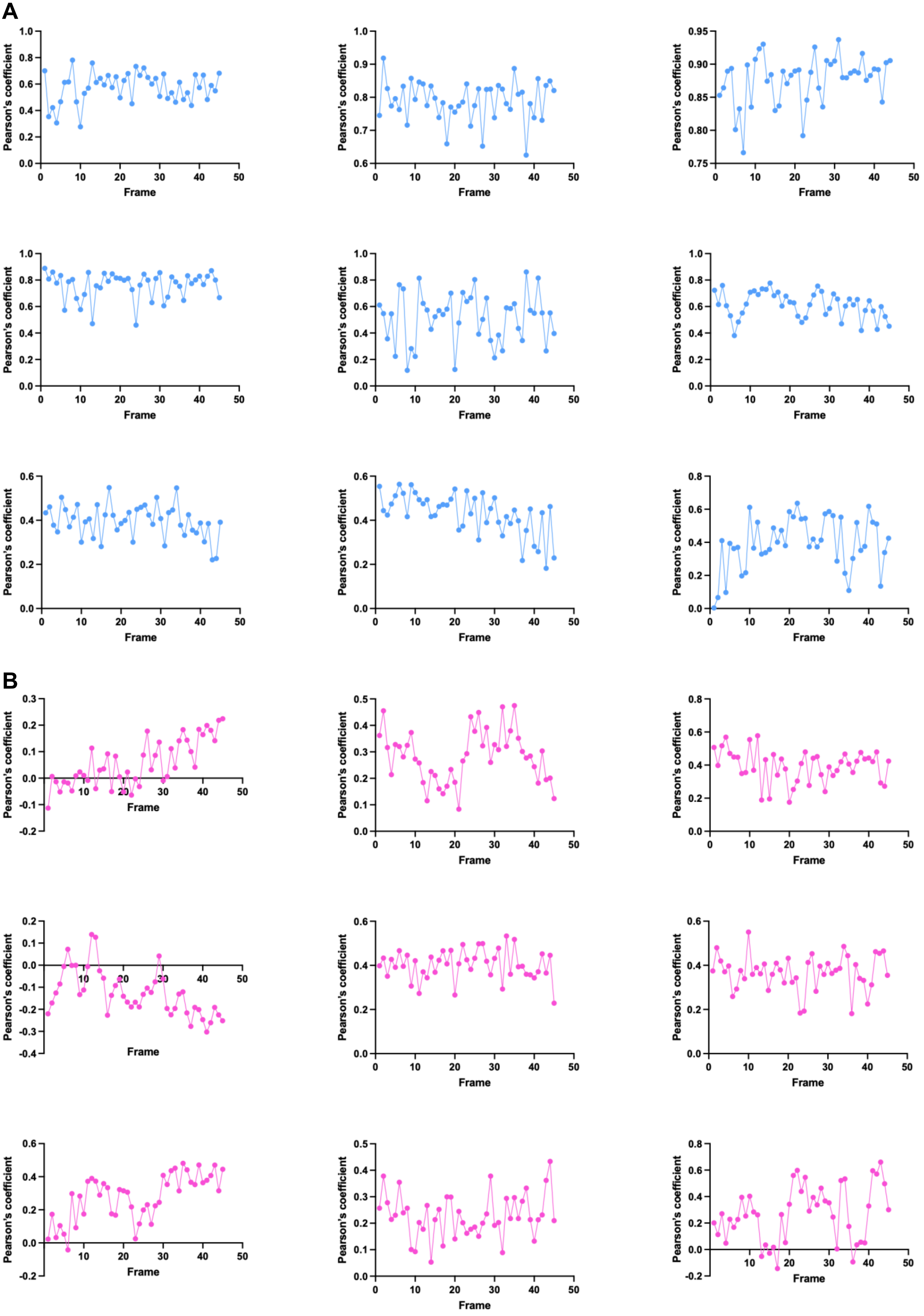
